# An in-silico analysis of experimental designs to study right ventricular function and pulmonary hypertension

**DOI:** 10.1101/2022.03.22.485347

**Authors:** M. J. Colebank, N.C. Chesler

## Abstract

*In-vivo* studies of pulmonary hypertension (PH) have provided key insight into the progression of the disease and right ventricular (RV) dysfunction. Additional *in-silico* experiments using multiscale computational models have provided further details into biventricular mechanics and hemodynamic function in the presence of PH, yet few have assessed whether model parameters are identifiable prior to data collection. Moreover, none have used modeling to devise synergistic experimental designs. To address this knowledge gap, we conduct an identifiability analysis of a multiscale cardiovascular model across four simulated experimental designs. We determine a set of parameters using a combination of Morris screening and local sensitivity analysis, and test for identifiability using profile likelihood based confidence intervals. We employ Markov chain Monte Carlo (MCMC) techniques to quantify parameter and model forecast uncertainty in the presence of noise corrupted data. Our results show that model calibration to only RV pressure suffers from identifiability issues and suffers from large forecast uncertainty in output space. In contrast, parameter and model forecast uncertainty is substantially reduced once additional left ventricular (LV) pressure and volume data is included. A comparison between single point systolic and diastolic LV data and continuous, time-dependent LV pressure volume data reveals that even basic, functional data from the LV remedies identifiability issues and provides substantial insight into biventricular interactions.

**Author Summary:** Computational models of cardiac dynamics are becoming increasingly useful in understanding the underlying mechanisms of disease. *In-silico* analyses are especially insightful in understanding PH and eventual RV dysfunction, as these conditions are diagnosed months to years after disease onset. Many researchers couple computational models with *in-vivo* experimental models of PH, yet few ever assess what data might be necessary or sufficient for parameter inference prior to designing their experiments. Here, we considered a multiscale computational model including sarcomere dynamics, biventricular interactions, and vascular hemodynamics, and assessed whether parameters could be inferred accurately given limited cardiac data. We utilized sensitivity analyses, profile likelihood confidence intervals, and MCMC to quantify parameter influence and uncertainty. We observed that RV pressure alone is not sufficient to infer the influential parameters in the model, whereas combined pressure and volume data in both the RV and LV reduced uncertainty in model parameters and in model forecasts. We conclude that synergistic PH studies utilizing computational modeling include these data to reduce issues with parameter identifiability and minimize uncertainty.

## INTRODUCTION

Computational modeling is a promising tool for understanding the origins as well as the progression of cardiovascular disease. Combined with invasive or non-invasive measurements, mathematical models of the cardiovascular system can forecast both the onset and worsening of disease (1–3). More recently, *multiscale* models that account for cardiovascular physiology across multiple spatial scales have been developed (4,5). The synergistic combination of *in-vivo* and *in-silico* methods have had notable success in understanding the progression of right ventricular (RV) failure in pulmonary hypertension (PH) (6–9). The left ventricle (LV) and septal wall (S) are highly coupled to RV function (10); hence, an impaired RV reduces biventricular energy efficiency and overall LV function (11). The use of mechanistic models and their physiologically based parameters can reveal additional details of PH progression, especially when combined with highly informative *in-vivo* data. However, these computational models suffer from numerous parameters and limited, noisy data available for parameter inference and model calibration (12). In these situations, a formal *identifiability analysis* can reveal which parameters to infer, and which data collection protocols are most informative for the model.

There are two main types of identifiability. Parameters are considered *structurally identifiable* if the model output, *f*(*t*;***θ***), is unique for every unique parameter set ***θ***. In addition to structural identifiability, parameters can also be *practically identifiable* if the parameters can be uniquely determined from limited and/or noisy data *y*(*t*). Structural identifiability assesses the model’s structure, and is determined using algebraic manipulations of the model (13–15) or by inferring parameters using noise-free, model generated data (1,16). Parameters that are deemed structurally identifiable can be assessed for practically identifiability in the presence of noisy and limited data. This type of analysis is imperative to inform *in-vivo* experimental designs, as model identifiability may dictate the frequency or quality of measurements necessary to calibrate the model.

Several authors have considered parameter identifiability in the context of cardiovascular modeling (1,15,17,18). Pironet et al. (15) pursued a structural identifiability analysis on a six-compartment model of the cardiovascular system. The study concluded that a combination of pressure and volume data was necessary to eliminate structural non-identifiability for the 13 parameters in their model. A follow up investigation by Pironet et al. used local sensitivity and profile likelihood analyses to conclude that, in a simplified three state model, four out of seven parameters were practically identifiable from swine data in the vena cava, aorta, and left ventricle (LV) (17). The studies by Colunga et al. (18) and Harrod et al. (1) used models including the LV, RV, and both the systemic and the pulmonary circulations. The former (18) used local sensitivity analysis and Markov chain Monte Carlo (MCMC) methods to deduce identifiable parameter subsets given limited data from heart transplant patients. The latter study (1) utilized similar sensitivity and MCMC techniques, and tested for structural identifiability by examining the marginal posterior distributions for each parameter after fitting the model to noise-free, model generated data from patients with PH due to left heart failure. Both studies showed that the entire set of model parameters could not be estimated with the available data, and instead deduced a smaller subset of model parameters that were both identifiable and physiologically meaningful.

These prior studies did not consider a multiscale model, nor did they account for biventricular interaction through the septal wall. This interaction is especially important during the progression of PH, as septal deformation can switch from rightward to leftward during chronic RV pressure overload (6,10). The cutting edge reduced order model of biventricular interaction is the three-segment (“TriSeg”) model developed by Lumens et al. (8). Two recent studies by van Osta (5,19) applied sensitivity analysis and uncertainty quantification methods to the TriSeg model, and identified which parameters were influential on model forecasts of RV, LV, and septal wall (S) strain. These investigations utilized non-invasive clinical data, whereas only a few studies have used the TriSeg model with *in-vivo* animal data (4,9,20,21). Animal models of PH provide novel insight into PH progression (22,23), yet it is unclear how informative *in-vivo* data from these experiments are for calibrating computational models.

To address these gaps in knowledge, this study investigates parameter identifiability for a multiscale model of biventricular interaction and cardiovascular dynamics. We utilize sensitivity analyses, the profile likelihood, and MCMC techniques to deduce practical identifiability of the model. We focus on data obtained from four experimental designs; three that are common for monitoring animal models of PH and focus on the RV (23–25), and an additional design that utilizes dynamic pressure-volume data in both the LV and RV (26). We generate both noise-free and noisy data from the model to test for parameter identifiability and analyze the output uncertainty in model simulations and several biomarkers of PH progression.

## MATERIALS AND METHODS

### Mathematical model

We consider a multiscale cardiovascular model describing sarcomere-level dynamics, biventricular interaction, and zero-dimensional (0D) hemodynamics. We summarize the mathematical model here and relegate individual component details to the Appendix.

The model consists of nine compartments: the systemic and pulmonary arteries and veins, the left and right atria, and a model accounting for interactions between the LV, RV, and S. A model schematic is provided in Fig 1.

**Fig 1.**
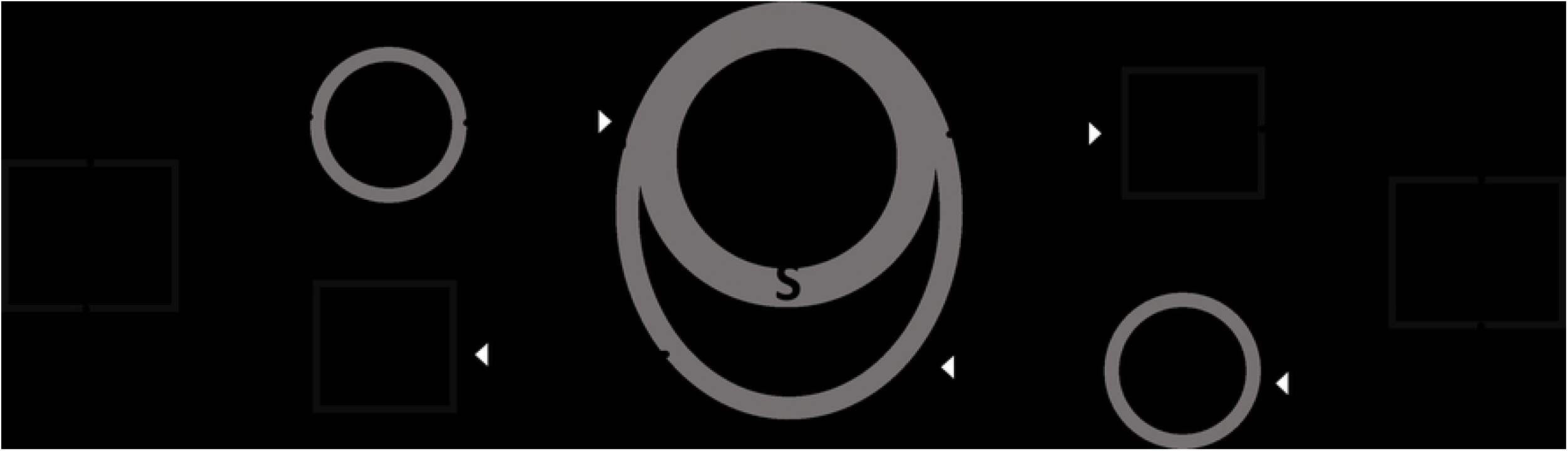
Model schematic. The computational model here consists of a lower order simulator of sarcomere dynamics within the left atrium (LA), left ventricle (LV), right atrium (RA), right ventricle (RV), and septum (S). The LV, RV, and S are simulated using the TriSeg model (8), and account for biventricular interaction. Lastly, a circuit model is used to describe the systemic arteries (SA) and veins (SV), as well as the pulmonary arteries (PA) and veins (PV).

### Sarcomere model

The sarcomeres in the atrial, ventricular, and septal walls are modeled as two passive elastic elements in parallel with an elastic and contractile element in series (27). The contractile sarcomere length, *L_sc_* (μm), and contractility, Γ (dimensionless), for each are dictated by ordinary differential equations (28). As described by Lumens et al. (8), changes in sarcomere length, *L_s_* (*μ*m), are dependent on myocardial strain, *ε_f_* (dimensionless), while changes in Γ depend on *L_sc_* and time *t* (s). Cardiac contractility is modeled as the sum of a rise and decay function, describing the binding of crossbridges, calcium fluctuations, and detachment of crossbridges during diastole. Active stress, *G_act_* (KPa), is determine as a function of *L_sc_* and Γ, whereas passive stress due to structural properties of the extracellular matrix, *G_ECM_* (KPa), and the giant protein Titin, *G_Titin_* (KPa), are strictly a function of sarcomere length. The total force generated from the sarcomere is then the sum of the active and passive forces

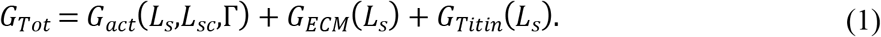

This subcomponent of the model constitutes a total of 27 parameters: 13 shared between the two atria, 13 shared between the LV, RV, and S, and a parameter describing the time delay of atrial contraction (see the Appendix).

### TriSeg Model

The sarcomere model is embedded within a cardiac tissue model of atrial dynamics and biventricular interaction (the “TriSeg” model (8)), and relates changes in blood volume *V*(*t*) (*μ*l) to myocardial strain *ε_f_*, using

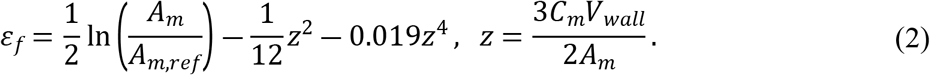

Here, *A_m_* (mm^2^) is the current mid-wall area of the chamber, *A_m,ref_* (mm^2^) is the reference mid-wall area, and *z* (dimensionless) is a curvature variable related to the ratio of wall volume, *V_wall_* (mm^2^), and radius of mid-wall curvature *C_m_* (mm^-1^) (8). Once *ε_f_* has been calculated and the corresponding *G_Tot_* is obtained from the sarcomere model, the mid-wall tension can be calculated as

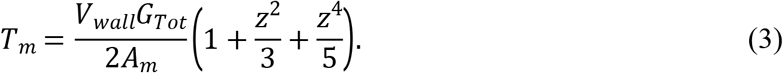

A balance in axial and radial tensions, *T_x_* and *T_y_* (see Appendix), is enforced

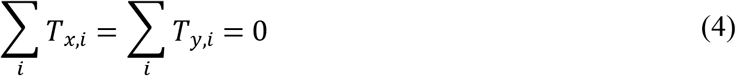

providing two differential algebraic equations (29). The cavity tensions are used to calculate the cavity pressures (see Appendix). In total, the cardiac chambers and TriSeg model contribute two algebraic constraints in equation (4), five wall volume parameters (*V_wall_*), and five reference area parameters (*A_m,ref_*).

### Hemodynamics model

The systemic and pulmonary arteries and veins are modeled as compliant compartments, with resistance elements between each compartment or cardiac chamber (18,30). In brief, changes in *V*, flow (*q*, *μ*l/s), and pressure *p* (KPa) are related via an electric circuit analogy

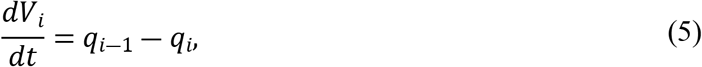

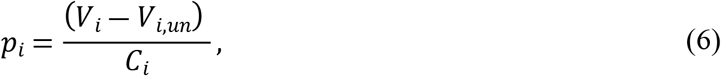

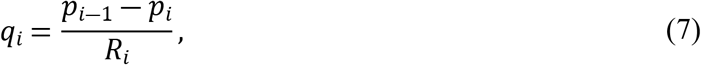

where *V_i,un_* (*μ*l) is the unstressed volume (see S1 text), *C_i_* (*μ*l KPa^-1^) is the vascular compliance, and *R_i_* (KPa s *μ*l^-1^) is the vascular resistance between compartments. Finally, we model the two atrioventricular valves (mitral and tricuspid), the two semilunar valves (aortic and pulmonic), and the resistor between the systemic veins and right atrium as diodes

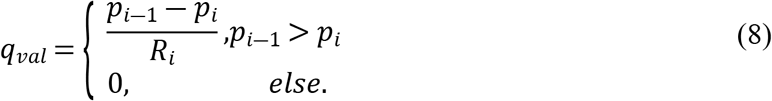

The hemodynamics model consists of eight differential equations for *V_i_*(*t*), eight resistance parameters, and four compliance parameters.

### Summary

The multiscale model consists of 18 differential equations (describing *L_sc_,* Γ, and *V*), two algebraic constraints (equation (4)), and a total of 49 parameters. Due to the algebraic constraints, the model constitutes a system of *differential algebraic equations* (DAEs) and is solved using the variable-step, variable-order ***ode15s*** solver available in MATLAB (Mathworks; Nantick, MA). Fig 2 shows nominal model predictions as well as the noise-corrupted data used in Bayesian parameter inference, discussed later.

**Fig 2.**
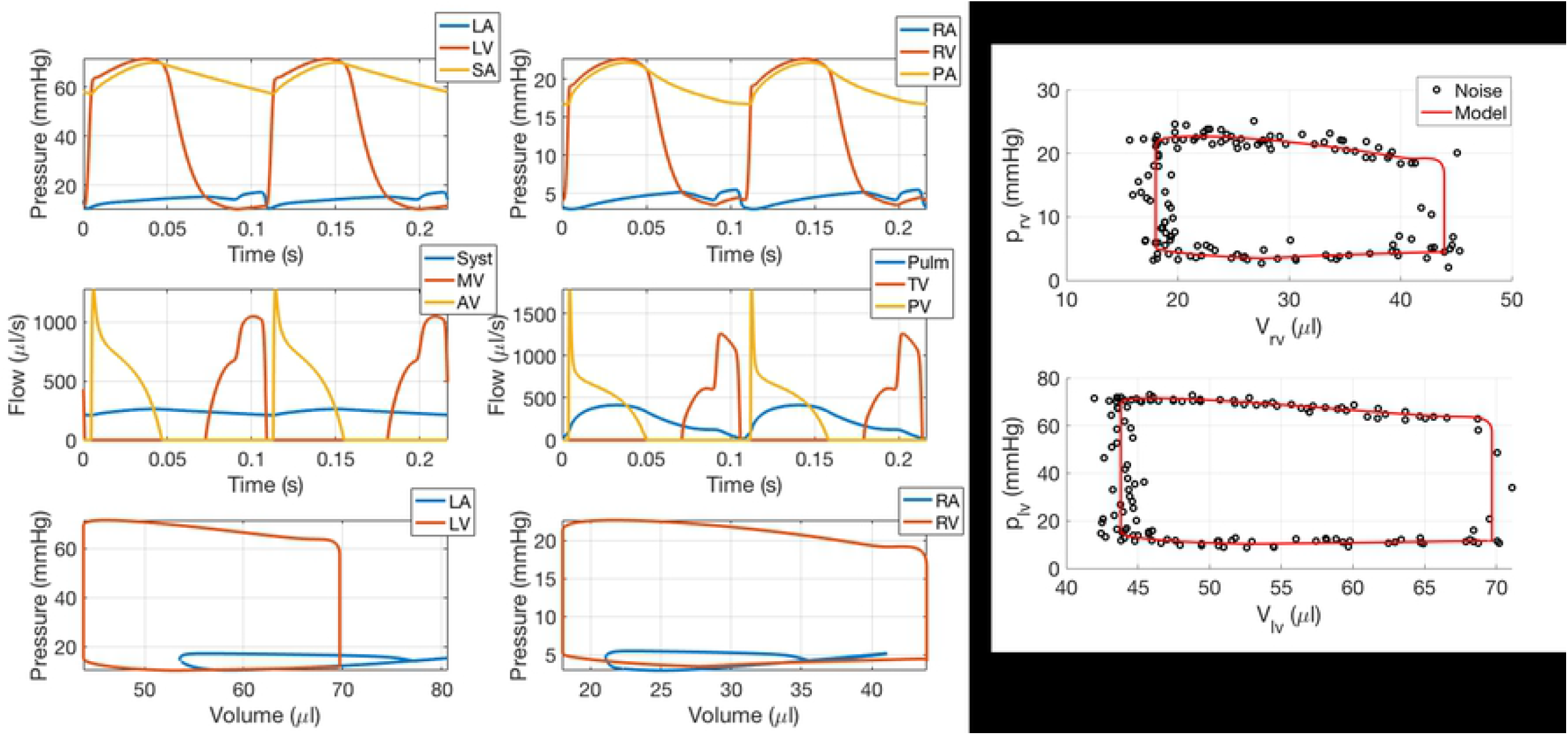
Nominal simulations and noise corrupted data. The nominal simulations are generated to match the data range reported by Philip et al. (23) in sham mice. Noise corrupted data is generated by adding additive, Gaussian errors with mean zero and a variance of 1.

#### Model sensitivity

Sensitivity analysis is an *a posteriori* identifiability method for determining which parameters are influential on a model output (15). *Local* sensitivity analysis perturbs parameters one at a time, and typically utilizes finite-difference approximations (30,31). In contrast, *global* sensitivity analysis samples parameters throughout the feasible parameter space, and includes variance based methods and screening methods (31,32). We utilize a Morris screening analysis (33) in combination with a local, derivative based sensitivity analysis for identifiability analysis. We utilize Morris screening over variance based methods since the model parameter space is large (***θ*** ∈ ℝ^49^). Prior studies have shown agreement between Morris’ indices and the total Sobol’ index (34), hence screening can be used to fix non-influential parameters.

The local sensitivity of a model output *f* with respect to a parameter, *θ_i_*, is approximated by the centered difference

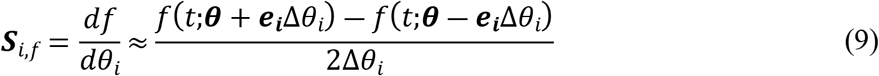

where *f*(*t*;***θ***) is the quantity of interest from the model, Δ*θ_i_* is the step change in parameter value, and ***e***_*i*_ is the *i*-th unit vector. For time-dependent outputs, we consider the 2-norm of the model output, i.e. 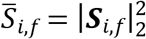. We account for differences in parameter magnitude by computing the log-scaled parameter sensitivity, *df/d*log(*θ*) (31,35).

The Morris’ screening approach computes the “elementary effects”

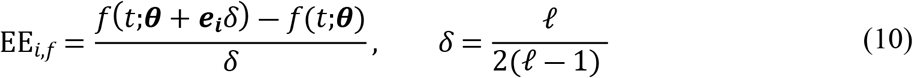

where *δ*(*ℓ*) is the parameter step size describing the “levels” of effects. Choosing *ℓ* to be even provides a more symmetric sampling distribution (33), hence we choose *ℓ* = 60 giving *δ* ≈ 0.51. Note that EE_*i,f*_ is a coarser approximation of model sensitivity than ***S***_*i,f*_, but is quantified over a larger parameter space. We scale parameters from their original value to the interval [0,1] as done previously (34), and utilize the algorithm provided by Smith (12) to construct our sampling methodology. The indices from the Morris method are

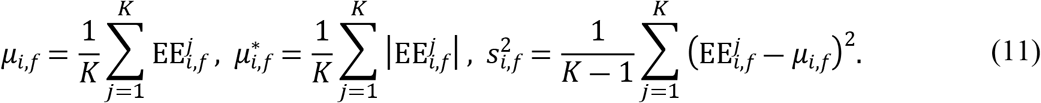

Here, *μ_i,f_* is the average of EE_*i,f*_, 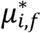 is an improved metric for average model sensitivity (34), and 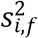 is the variance of EE_*i,f*_. We use the combined index, 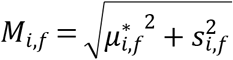, to measure a parameter’s influence (36).

Small values of either the local sensitivity index 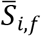 or the screening index *M_i,f_* indicate that a parameter is *non-influential,* i.e. it has minimal effect on *f.* As discussed next, these indices assess whether a model parameter is identifiable.

#### Practical parameter identifiability

In this work, we assess parameter identifiability using three techniques. The first is through the local and global sensitivity metrics discussed above. Next, we consider the profile likelihood, which provides information about whether each *θ_i_* is identifiable from a given set of data. Lastly, we use MCMC methods for Bayesian inference, and utilize the marginal posterior distributions to assess parameter identifiability.

### Sensitivity based identifiability

Parameters that have little effect on the model output are considered non-identifiable, since they do not affect the quantity of interest (12), and should be fixed before conducting inference. We employ a two-part parameter fixing methodology using the results from Morris screening and local sensitivity analysis.

A parameter is deemed *non-influential* for all outputs *f* if its index *M_i,f_* is less than the average 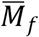 for all parameters *i* = 1,2,...,*P*

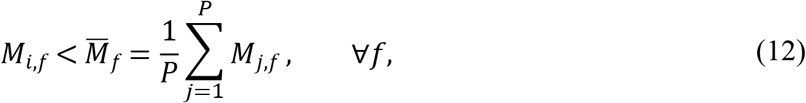

where *f* is one of the model outputs (5,32). Parameters that are less than this threshold for all outputs are considered non-influential for inference and are fixed.

After using the Morris screening approach, the subset is analyzed by conducting a local sensitivity analysis around the nominal parameter values. The Fisher information matrix, 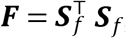, must be non-singular for gradient based parameter estimation, hence its utility in parameter identifiability (37). If ***F*** is invertible but has a large condition number, then some of the sensitivities are nearly linearly dependent and the subset requires further reduction. We use an eigenvalue-eigenvector analysis method to determine which parameters cause the ill-conditioning of ***F*** (14,38), and fix these parameters at their nominal value.

### Profile likelihood

The most common and robust technique for assessing practical identifiability is the profile likelihood (13,15). This technique increments a fixed parameter, *θ_i_*, while minimizing the negative log-likelihood for all other parameters in the subset, i.e.

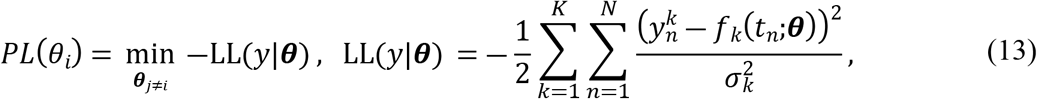

where *y^k^* is the *k*-th data source, *f_k_* is the corresponding model output, LL(*y*|***θ***) is the log-likelihood, 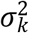 is the noise variance for the data source, and *N* is the number of data points. The corresponding profile likelihood confidence intervals for *θ_i_* are (13)

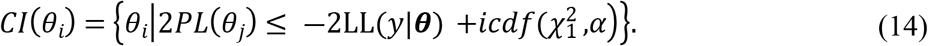

Each *CI*(*θ_i_*) is constructed around the optimal estimate, ***θ***\*, and depends on the inverse cumulative distribution function of the chi-squared distribution, 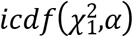, with one-degree of freedom and confidence level *α* (13). If *PL*(*θ_i_*) is completely flat (e.g., *CI*(*θ_i_*) is infinite), then *θ_i_* is deemed *structurally non-identifiable* and cannot be uniquely determined due to model structure. If only one side of *PL*(*θ_i_*) is flat, then *θ_i_* is considered *practically non-identifiable,* and could become identifiable if more data was available for inference (16).

### Bayesian inference

Bayesian parameter inference using MCMC is more computationally expensive than gradient based inference, but provides detailed insight into parameter relationships and avoids local minima in the likelihood (39–41). We use the DRAM algorithm (42), which is described in depth elsewhere (31,43). In short, the goal of MCMC is to approximate the posterior distribution

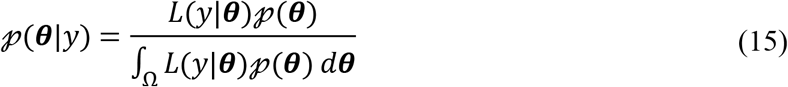

where 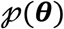 is the prior distribution, *L*(*y*|***θ***) is the likelihood, and the denominator of equation (15) is a normalization factor, approximated using Monte Carlo sampling. Model parameters are sampled from a proposal distribution to compute the likelihood *L*(***θ***\*|*y*), where ***θ***\* is the proposed parameter values. The proposed parameter vector is accepted if the ratio of the likelihood values between ***θ***\* and the previous value of ***θ*** are greater than some random realization from a unit normal distribution. To reduce parameter stagnation or random-walk behavior, a second proposal parameter set is generated from a narrower distribution if ***θ***\* is rejected (43). The DRAM algorithm updates the covariance matrix of the proposal after sequential adaption intervals, improving the proposed values of ***θ***\* (43).

We utilize DRAM on a set of noisy data, generated by the model and corrupted with noise. To ensure adequate parameter space coverage, we first initialize a gradient based optimization from twelve randomized initial parameter sets and minimize the residual sum of squared errors for the given experimental conditions (defined in the next section). The optimal parameter vector, ***θ**_SSE_,* is used as a starting value for DRAM and the Hessian matrix obtained from the optimization is used as the initial covariance matrix to preserve possible sampling asymmetry (12). We implement this using the freely available DRAM package developed Haario et al. (42) in MATLAB. In situations where the model is unstable or crashes, we return a large value for the residual sum of squares (39). We assess parameter identifiability by visualizing the marginal posterior densities 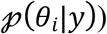); longer, unbounded tails in the posterior suggest issues with parameter identifiability.

#### Simulated experiments and additional outputs

Several experimental designs are commonly used for *in-vivo* PH studies (23,24). We are interested in using the computational model to infer parameters indicative of heart function; hence, we consider different assortments of ventricular pressure and volume data. Our experimental designs are:

1. Dynamic measurements of RV pressure (***f***_1_ = [*p_RV_*(*t*)]);
2. Dynamic measurements of RV pressure and volume (***f***_2_ = [*p_RV_*(*t*),*V_RV_*(*t*)]);
3. RV pressure and volume measurements, as well as systolic and diastolic pressure and volume in the LV 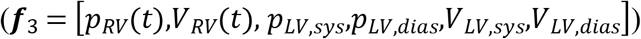; and
4. Dynamic measurements in both the RV and LV (***f***_4_ = [*p_RV_*(*t*),*V_RV_*(*t*), *p_LV_*(*t*),*V_LV_*(*t*)]).

The first two scenarios correspond to *in-vivo* recordings from pressure (24) or pressure-volume catheters (3,23). The third includes additional information on the LV obtained by echocardiography (23). Finally, the fourth experimental design represents a realistic, but underutilized, scenario that includes pressure-volume measurements in both the RV and LV (11,26).

We perform all sensitivity and identifiability analyses with respect to the pressure and volume forecasts considered in the four experimental designs above. Noisy pressure and volume data are generated by adding zero mean, white Gaussian noise, with a variance of 1. Parameter subsets for each experimental design are contrasted, with a common subset determined across all designs. To better understand the consequences of limited data, we construct profile likelihood confidence intervals and analyze the parameter posterior distributions for each design. In the latter case, we compare the maximum *aposteriori* estimates with the known, data generating parameters. Lastly, we propagate uncertainties in the model parameters to simulated outputs via the posterior distributuons. This includes LV, RV, and S engineering strain (19) as well as mean pulmonary artery pressure, RV stroke volume (the difference between end-diastolic and end-systolic volumes), pulmonary arterial elastance (the difference in mean pulmonary artery and left atrial pressure over stroke volume), RV end-systolic elastance, and RV ventricular-vascular coupling (23). The source code of the mathematical model and relevant analyses can be found at https://github.com/mjcolebank/Colebank_Identifiability_2022.

## RESULTS

Before beginning the sensitivity analysis, several parameters were excluded for physiological reasons. For instance, the reference length of the sarcomeres, *L_s,ref_* were excluded from analysis since these values are consistent across experimental designs. Table 1 summarizes the model parameters that are considered in our analyses. A detailed description of how parameter values are calculated can be found in the S1 text.

**Table 1.** Parameters, their description, and information regarding the sensitivity analyses.

| Parameter | Description | Used in sensitivity analyses | Parameter | Description | Used in sensitivity analyses |
| --- | --- | --- | --- | --- | --- |
| $V_{LA,wall}$ | LA wall volume | X | $\tau_{offset,A}$ | Offset of atrial systole | |
| $V_{LV,wall}$ | LV wall volume | X | $\bar{\sigma}_{act,A}/\bar{\sigma}_{act,V}$ | Active stress scaling | X |
| $V_{RA,wall}$ | RA wall volume | X | $\bar{\sigma}_{pas,A}/\bar{\sigma}_{pas,V}$ | Passive stress scaling | X |
| $V_{RV,wall}$ | RV wall volume | X | $L_{s,pas,ref,A}/L_{s,pas,ref,V}$ | Reference length for passive wall constituents | |
| $V_{S,wall}$ | S wall volume | X | $\beta_{pas,A}/\beta_{pas,V}$ | Stiffness of passive element | X |
| $A_{m,ref,LA}$ | LA reference area | X | $k_{1,A}/k_{1,V}$ | Nonlinear scaling of Titin stiffness | X |
| $A_{m,ref,LV}$ | LV reference area | X | $R_{a,val}$ | Aortic valve resistance | X |
| $A_{m,ref,RA}$ | RA reference area | X | $R_{m,val}$ | Mitral valve resistance | X |
| $A_{m,ref,RV}$ | RV reference area | X | $R_{p,val}$ | Pulmonic valve resistance | X |
| $A_{m,ref,S}$ | S reference area | X | $R_{t,val}$ | Tricuspid valve resistance | X |
| $L_{s,ref,A}/L_{s,ref,V}$ | Reference sarcomere length at zero strain | | $R_{vc}$ | Vena Cava resistance | X |
| $L_{s,iso,A}/L_{s,iso,V}$ | Elastic series element length in isometric state | | $R_{pv}$ | Pulmonary venous resistance | X |
| $v_{0,A}/v_{0,V}$ | Velocity of sarcomere shortening | X | $R_{sys}$ | Systemic circulation resistance | X |
| $L_{sc,0,A}/L_{sc,0,V}$ | Contractile element length | | $R_{pulm}$ | Pulmonary circulation resistance | X |
| $\Gamma_{rest,A}/\Gamma_{rest,V}$ | Resting contractility | | $C_{sa}$ | Compliance of systemic arteries | X |
| $\tau_{rise,A}/\tau_{rise,V}$ | Rise in contractility scaling | X | $C_{sv}$ | Compliance of systemic veins | X |
| $\tau_{decay,A}/\tau_{decay,V}$ | Decay in contractility scaling | X | $C_{pa}$ | Compliance of pulmonary arteries | X |
| $\tau_{sys,A}/\tau_{sys,V}$ | Length of systole | X | $C_{pv}$ | Compliance of pulmonary veins | X |

We ran the Morris screening algorithm using 100 randomized initializations. Fig 3 shows the parameter ranking *M_i,f_* using the mean effect *μ\** and corresponding variance *s*^2^ for the RV and LV pressures and volumes. See S1 text for individual results from the Morris screening analysis as well as parameter bounds for sampling. Sensitivity results were analyzed by comparing the parameter ranking *M_i,f_* to the mean effect 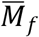 for each ventricular pressure and volume. All four compliances were consistently ranked within the most influential parameters, while other parameters describing cardiac chamber dynamics (e.g., *A_m,ref_*) varied with the output. We fixed parameters that were less influential than 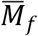 for all four outputs (i.e., pressure and volume in the RV and LV). This reduced our parameter subset from 38 to 17 parameters, shown in Table 2.

**Fig 3.**
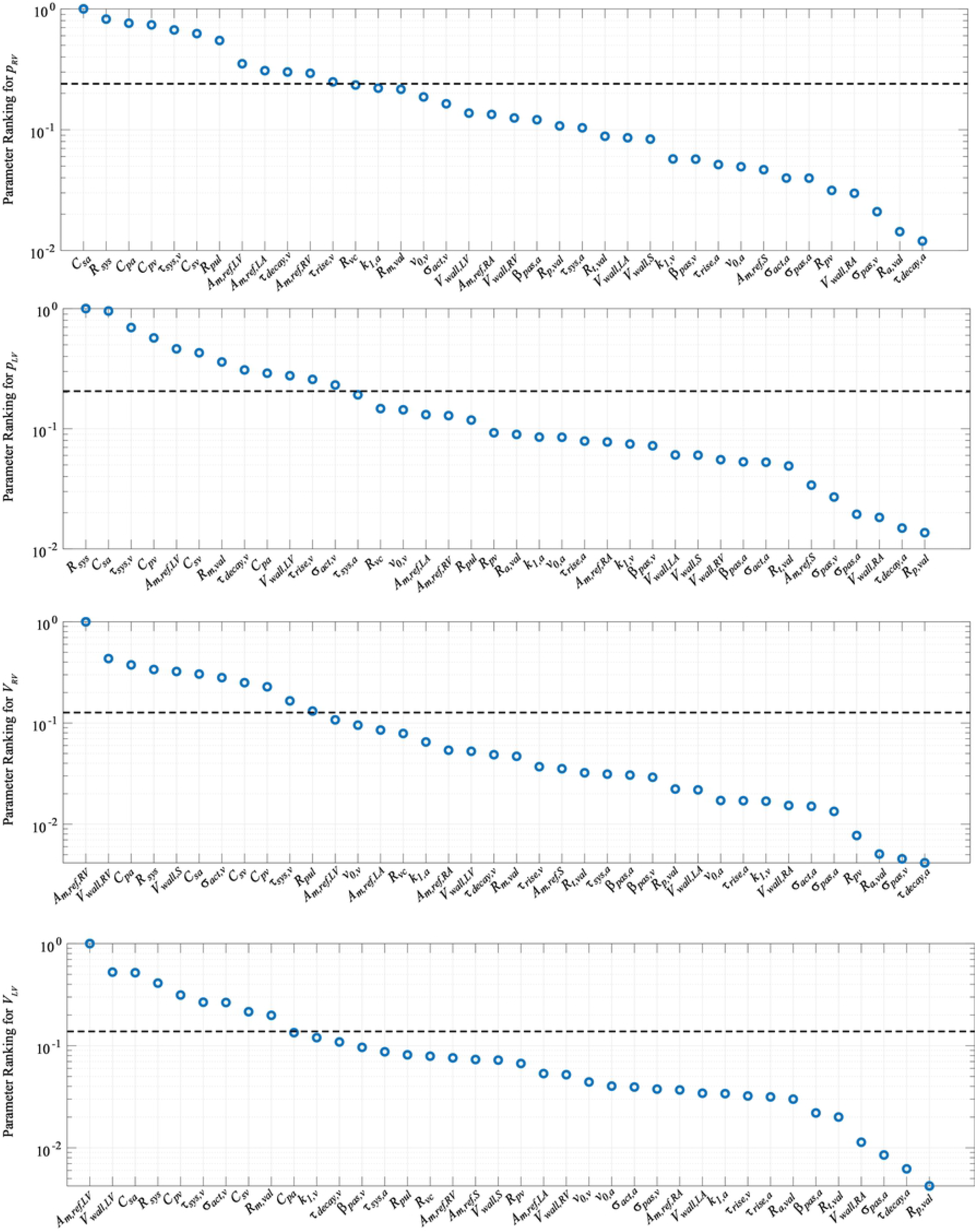
Sensitivity results from the Morris screening algorithm. Parameter ranking is based off of the index 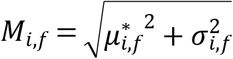. The dotted line in each plot denotes the average model sensitivity for each output.

**Table 2.**
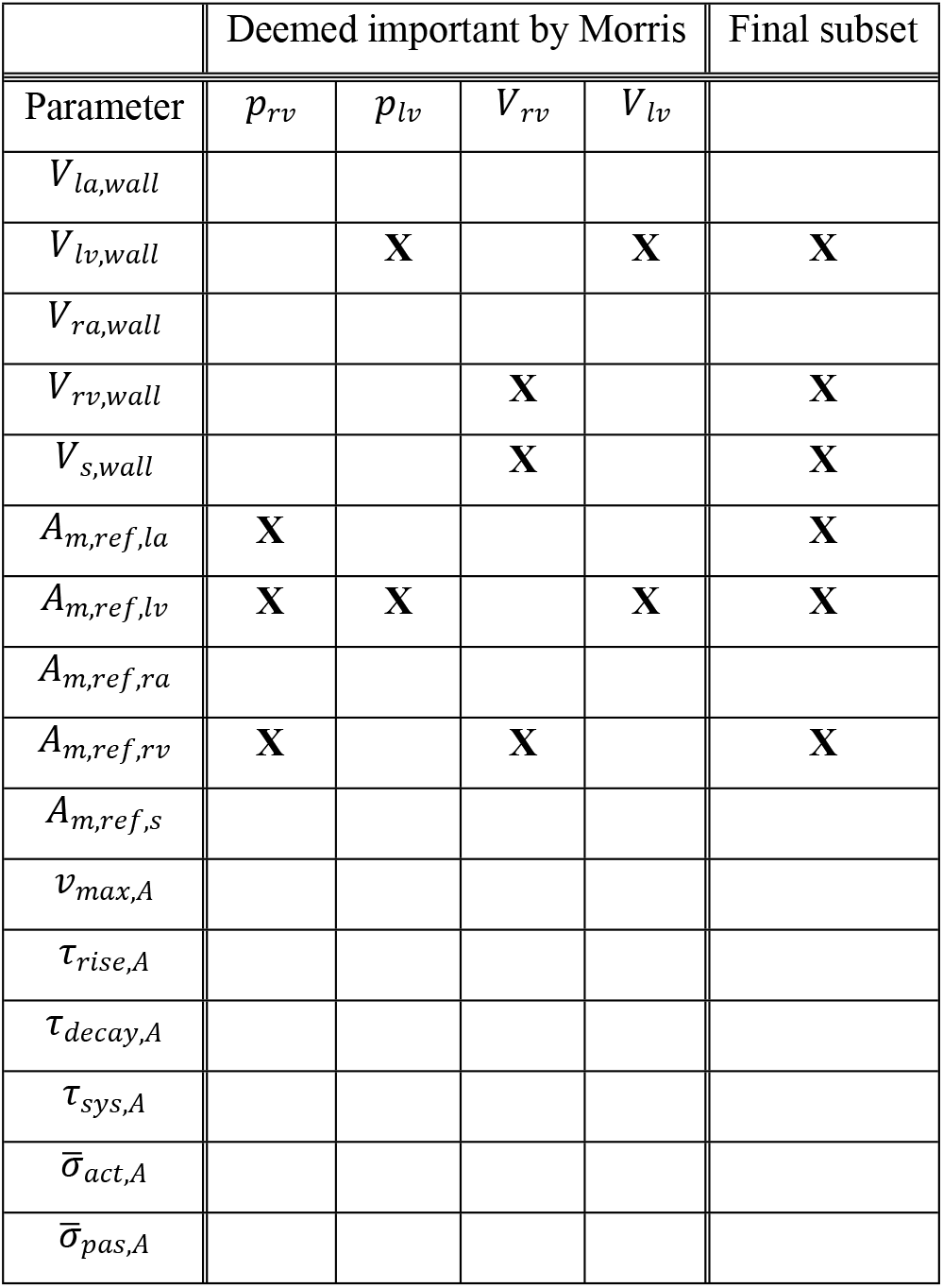
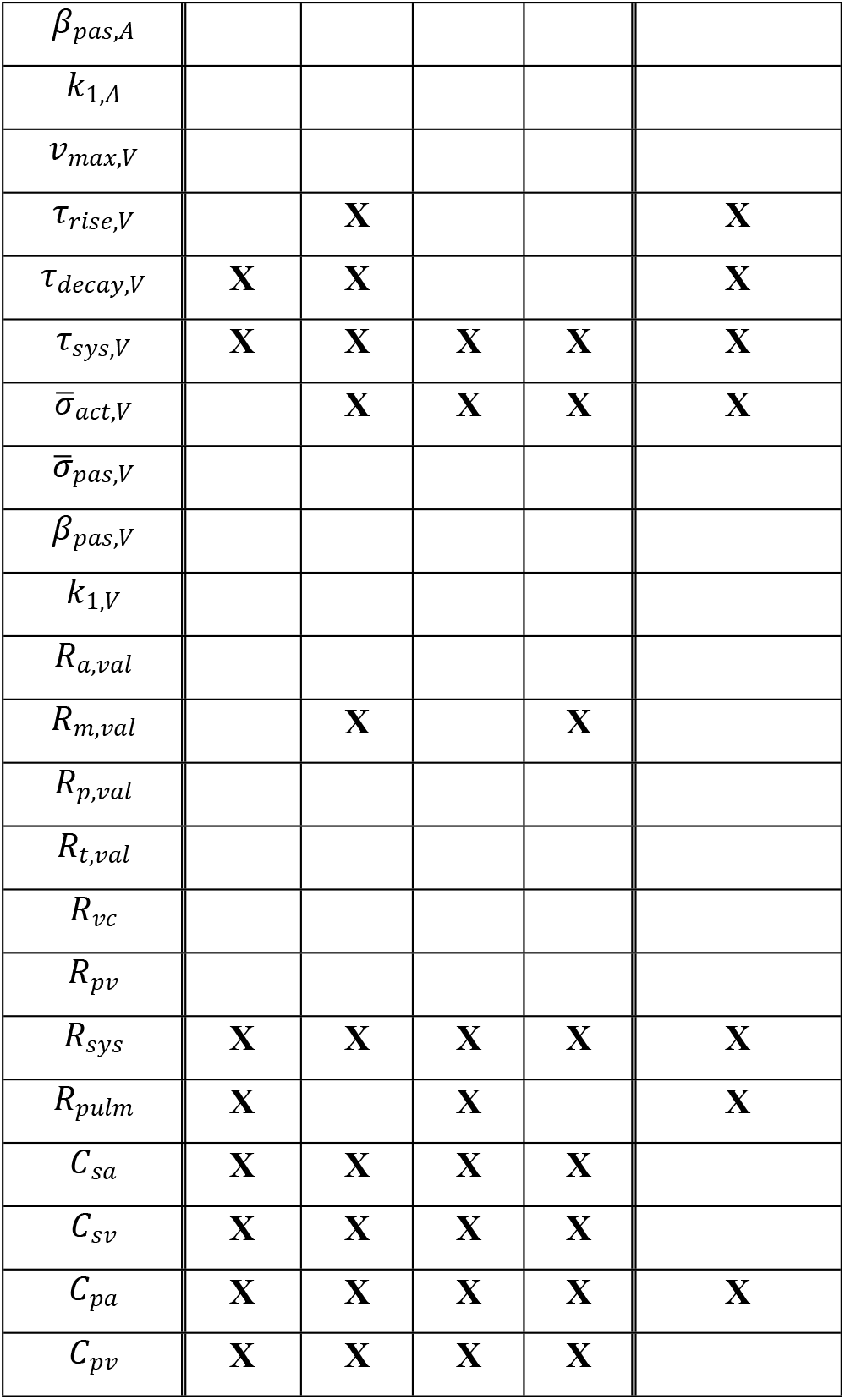
Parameters deemed influential by Morris screening and included in the final subset after using a local sensitivity based identifiability analysis.

We conducted a local sensitivity analysis on the reduced subset of 17 parameters using the designs ***f***_1_, ***f***_2_, ***f***_3_, and ***f***_4_ as the quantity of interest. The local sensitivity of these designs with respect to the 17 parameters are used to construct the Fisher information matrix, ***F***. Using the SVD decomposition, we reduced the parameter subset until cond(***F***) ≤ 10^8^ for each design, providing a subset of 13 parameters deemed identifiable for all four designs. Parameters fixed by the SVD method included mitral valve resistance, *R_m,val_*, and compliance in the systemic arteries, systemic veins, and pulmonary veins (*C_sa_, C_sv_,* and *C_pv_,* respectively). This final subset, shown in Table 2, was used in the profile likelihood and MCMC analysis.

Profile likelihood based confidence intervals are constructed using the noise-free, model generated data. We construct the confidence intervals ± 50% away from the true parameter value, with the confidence level cutoff for each design calculated using equation (14) with an *α* = 0.95 confidence level. The profile likelihood results, displayed in Fig 4, show that only the last experimental designs, ***f***_4_, provided finite confidence bounds for all 13 parameters. Sharp edges in the profile likelihood correspond to local minima and/or incompatible parameter sets corresponding to a failure in the DAE solver. The parameters *A_m,ref,RV_, τ_rise,V_, τ_decay,V_, τ_sys,V_,R_sys_*, and *C_pa_* were identifiable for all four experimental designs. The remaining seven parameters (*V_wall,LV_, V_wall,RV_, V_wall,S_, A_m,ref,LA_, A_m,ref,LV_ σ_act,V_,* and *R_pulm_*) varied in their identifiability with each experimental design.

**Fig 4.**
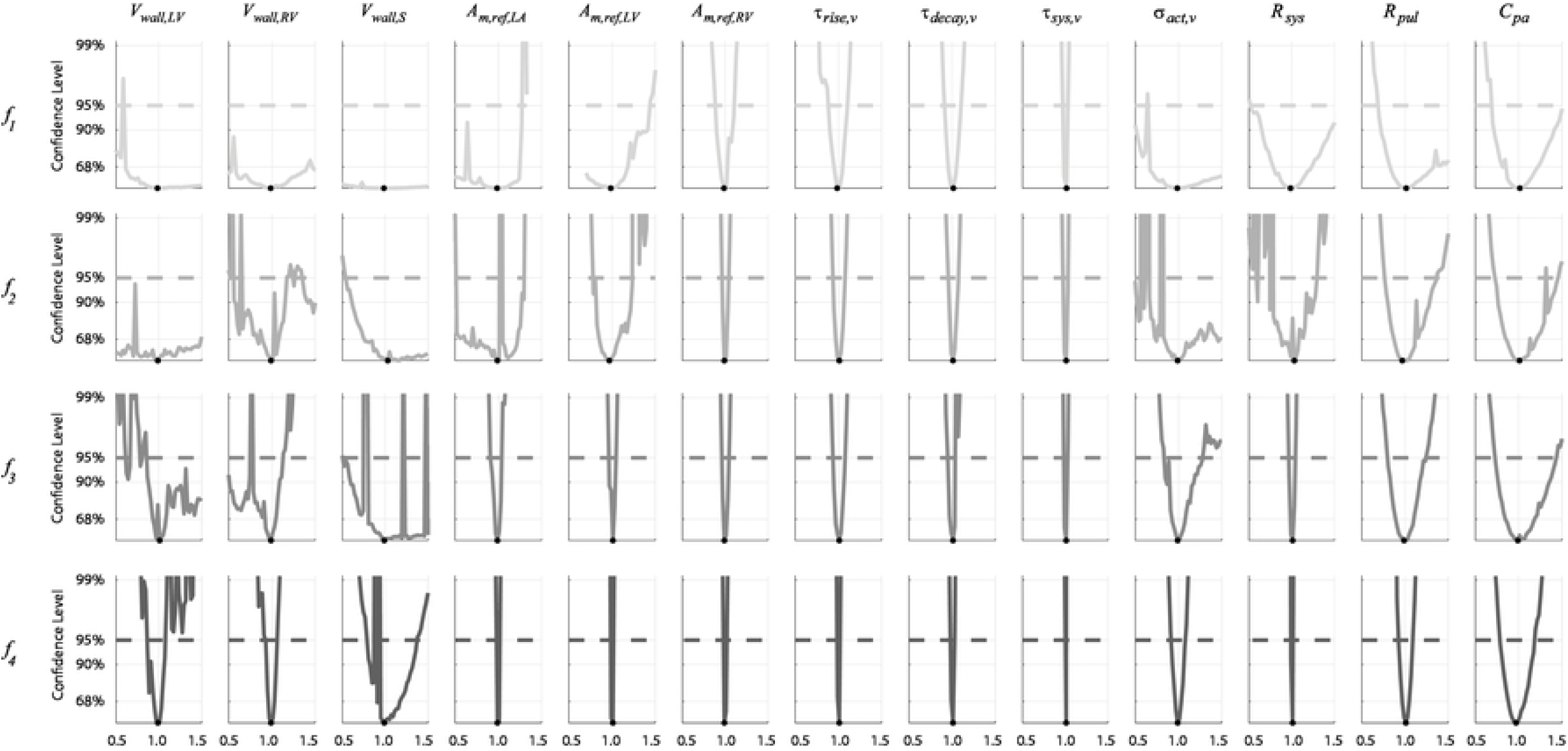
Profile likelihood confidence intervals. Confidence intervals are constructed by fixing one parameter and inferring all others over a range of values. Each row corresponds to a different experimental design. Note that the minimally informative experimental designs (***f***_1_ and ***f***_2_) have non-identifiable parameters, indicated by infinite or one-sided confidence bounds. In contrast, inclusion of LV data (***f***_3_ and ***f***_4_) remedy the issue of non-identifiable parameters in the set. Large deviations in the profile likelihood correspond to local minima and parameter sets that are incompatible for the system of DAE’s.

Noise corrupted data generated by the model is used in the likelihood defined in equation (13). We use minimally informative priors (i.e., with a large variance) for each parameter and initialize the DRAM algorithm using the optimal parameter vector ***θ***_*SSE*_ and estimated covariance matrix from twelve randomly selected initial guesses. MCMC is run for 50,000 iterations, with the initial 10,000 being left out as a “burn-in” period. We separate the results from MCMC into three groups: parameters representing the heart chambers’ geometry (Fig 5), parameters within the sarcomere model (Fig 6), and hemodynamic parameters in the circulatory model (Fig 7). Fig 5, Fig 6, and Fig 7 show three of the twelve MCMC chains as well as the posterior distribution calculated using kernel density estimation. The posterior distributions are relatively wide when only using RV pressure data (***f***_1_), but additional data in the subsequent experimental reduce the posterior widths. All the marginal posterior distributions contain the true, data generating parameters, though some of the posteriors’ modes are unaligned with the true parameters. Additional pairwise plots, provided in S2 text, suggest some correlation between variables. One chain using ***f***_3_ shows tight, narrow correlations, likely due to a poor initialization during the optimization and inadequate exploration of the parameter space. However, the other eleven instances suggest minimal correlations between variables for ***f***_3_. In general, 50,000 iterations appear sufficient for most of the MCMC results; however, the addition of static LV data with ***f***_3_ causes some suboptimal mixing for the parameter *σ_act,v_*. The MCMC chains appear to converge quicker when using the most detailed experimental design, ***f***_4_. The median acceptance rates for the twelve chains are 29.8%, 21.7%, 36.1%, and 49.0% for designs ***f***_1_, ***f***_2_, ***f***_3_ and ***f***_4_, respectively.

**Fig 5.**
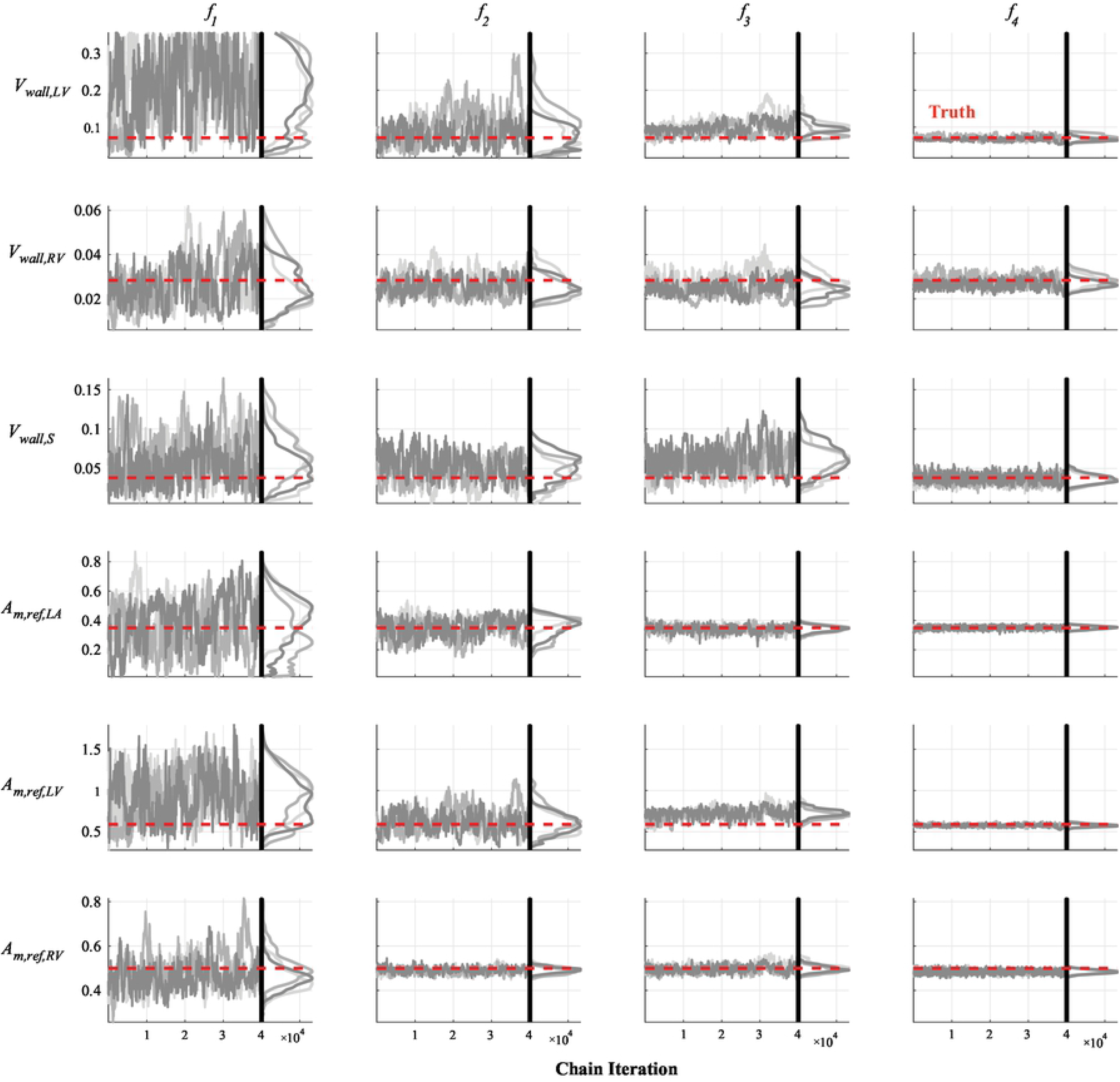
Chain iterations and marginal posteriors after MCMC for the TriSeg parameters. The model parameters indicative of the TriSeg geometry (wall volume, *V_wall_*, and reference mid-wall area, *A_m,ref_*) are shown for each experimental design, corresponding to each column. The true, data generating parameters corresponding to the outputs in Fig. 2 are shown as red lines. Three of the twelve initializations of MCMC are shown in different shades of gray. The marginal posterior distributions for the simplest experimental design (***f***_1_) are much wider than the subsequent more informed experimental designs, suggesting an improvement in practical identifiability.

**Fig 6.**
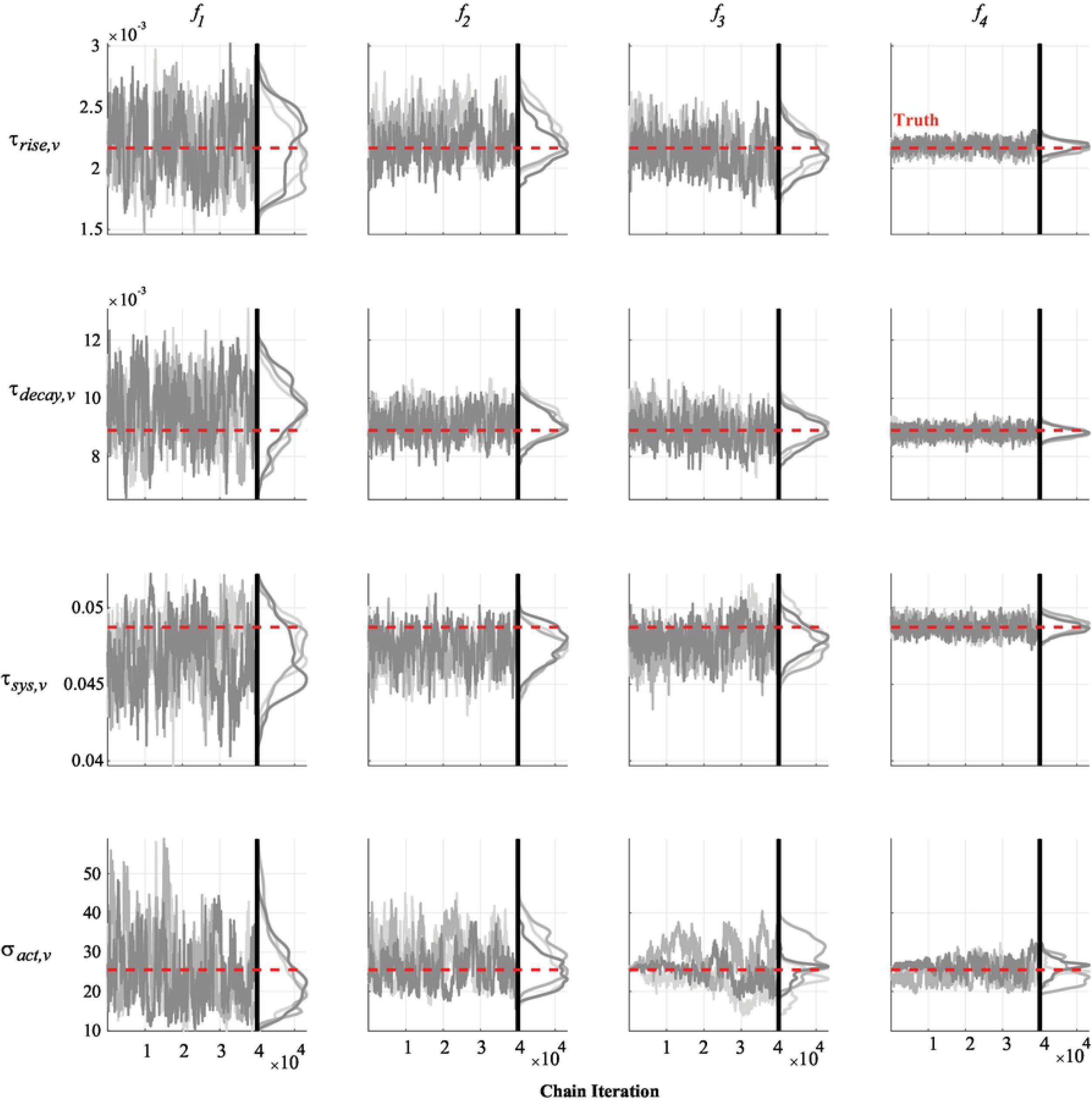
Chain iterations and marginal posteriors after MCMC for the sarcomere parameters. Similar to Fig. 5, three of the twelve MCMC instances are provided for the sarcomere parameters important for the rise, decay, and length of fiber shortening (*τ_rise,v_, τ_decay,v_*, and *τ_sys,v_*, respectively), and maximal active force generation (*σ_act,v_*). Note that all four experimental designs (given by each column) provide sufficient information to the likelihood so that the true data generating parameters (in red) are within the marginal posteriors.

**Fig 7.**
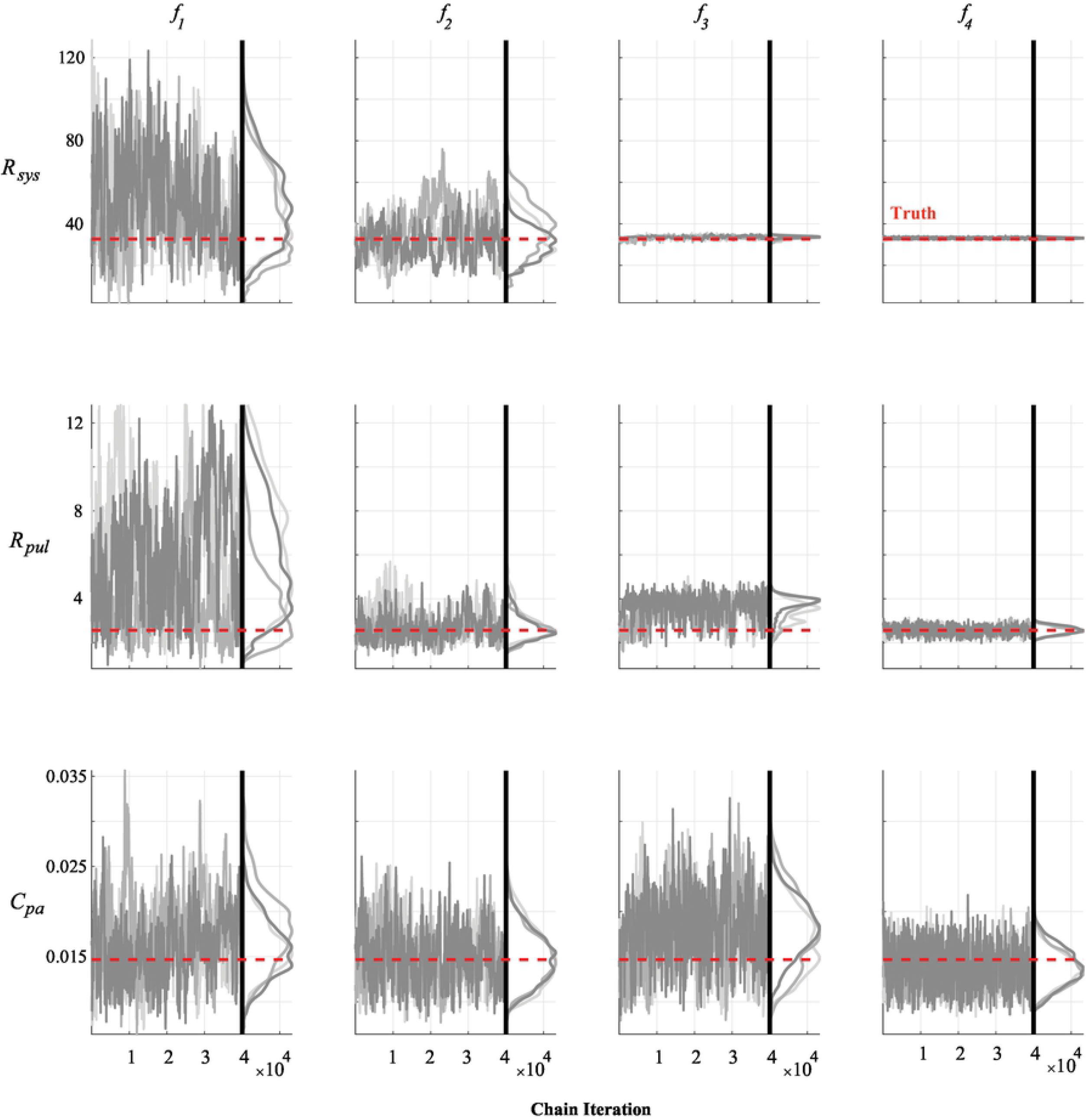
Chain iterations and marginal posteriors after MCMC for the hemodynamic compartment parameters. As in Figs 5 and 6, three of the twelve MCMC instances are provided for systemic vascular resistance, *R_sys_,* pulmonary vascular resistance, *R_pul_*, and pulmonary arterial compliance, *C_pa_.* Though the marginal posteriors do contain the true parameters (in red) within the marginal posteriors for the simplest design (***f***_1_, first column), additional data in the other designs substantially reduce posterior uncertainty.

We propagate the uncertainties in model parameters to the outputs by subsampling from the posterior distributions. To account for any across chain variation, we draw fifty samples from the twelve different MCMC instances, giving 600 realizations from the posteriors. Fig 8 displays the noise-corrupted data, average response from the agglomerated samples, and one standard deviation from the average response. The results from the initial design, ***f***_1_, show little uncertainty in RV pressure, but large uncertainty in forecasts of LV pressure and both chamber volumes. In contrast, ***f***_2_, ***f***_3_, and ***f***_4_ show reduced uncertainty once more data is added to the likelihood function. Note that the addition of dynamic LV pressure and volume in ***f***_4_ had relatively minimal effects on uncertainty when compared to only including systolic and diastolic values with ***f***_3_. We recast these results into pressure-volume loops in Fig 9 and provide the 600 realizations in addition to the agglomerated average and the true model simulations. Even in the absence of atrial pressure or volume data, additional volume measurements in both the LV and RV reduce the uncertainty of atrial dynamics. LV data reduces the uncertainty substantially in ***f***_3_ and ***f***_4_ as shown previously in Fig 8.

**Fig 8.**
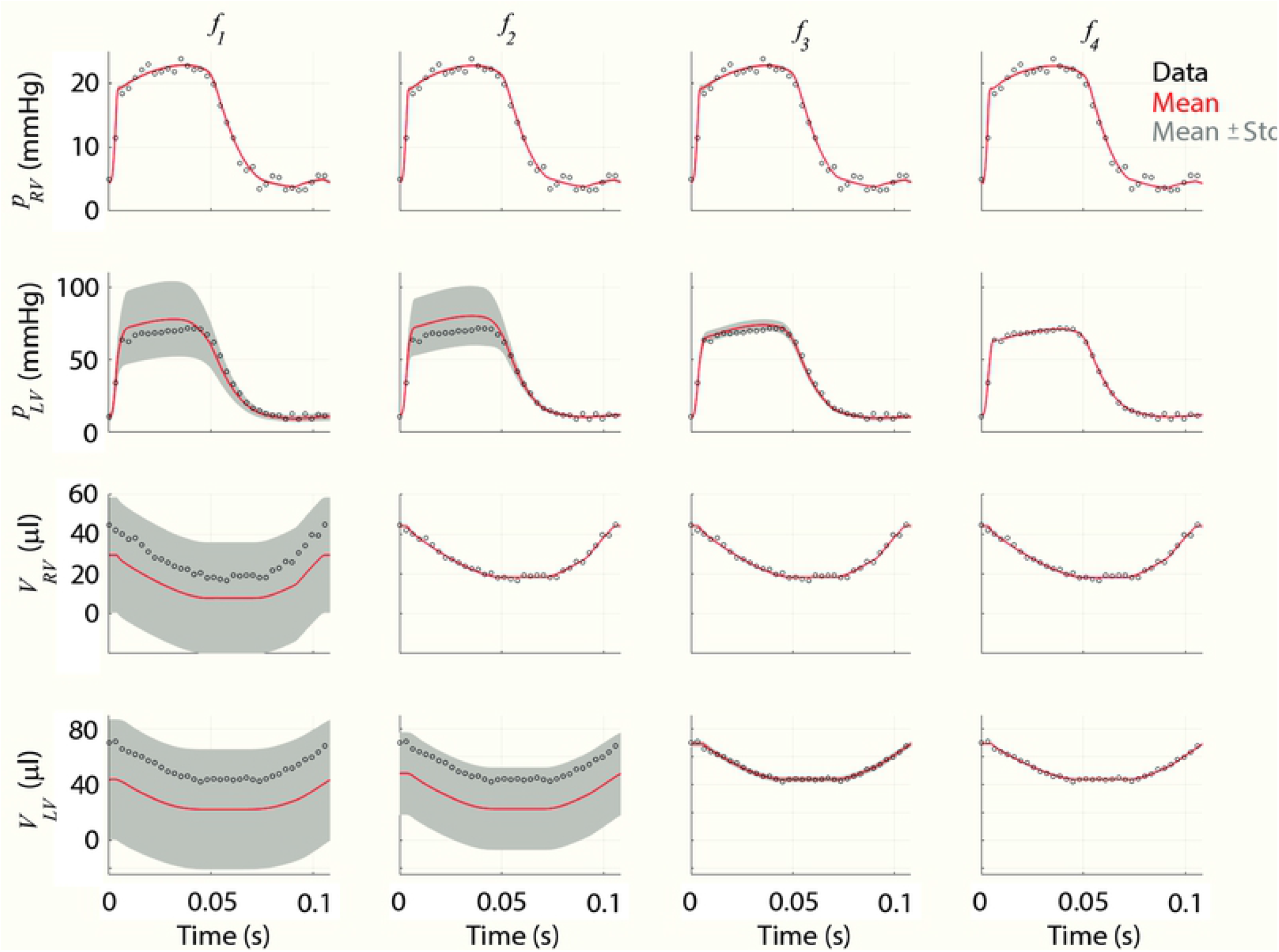
Output uncertainty in RV and LV pressures and volumes for each experimental design. The average model response (red) as well as ± one standard deviation (Std., gray) are provided along with the data (black circles) for each experimental design, corresponding to each column. In the first design, ***f***_1_, only RV pressure is used in the likelihood, hence the uncertainty in RV volume and LV forecasts are substantially larger than that of the RV pressure. As more data is included, uncertainty in model forecasts is reduced. Note that differences between *f*_3_ and *f*_4_ are less pronounced.

**Fig 9.**
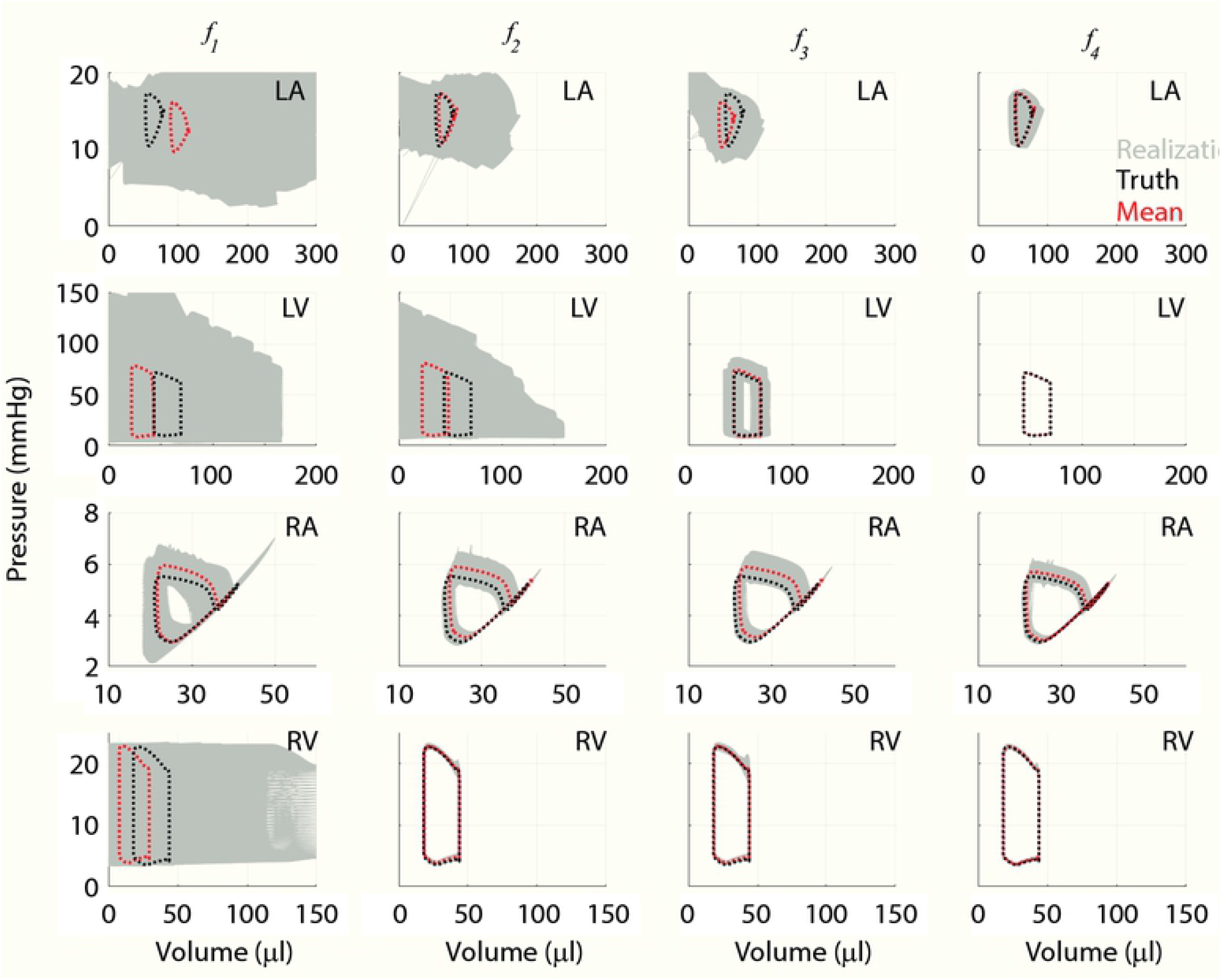
Output uncertainty in cardiac pressure-volume loops. Realizations in forecasts of chamber PV loops in the LA (first row), LV (second row), RA (third row), and RV (fourth row). The simplest design (***f***_1_) has the largest uncertainty in simulated PV loops, except for RV pressure, which is accounted for in the likelihood. Subsequent experimental designs substantially reduce uncertainty bounds in the RV and RA (***f***_2_) and eventually in the LV and LA (***f***_3_ and ***f***_4_).

In addition to outputs that are linked to the collected data, we investigate the uncertainty in ventricular wall strain and outcomes typically quantified during *in-vivo* PH studies. Engineering strain for the LV, RV, and S walls are provided in Fig 10. Strains are bounded between 5% and −20%, and there was a reduction in uncertainty when additional data was included in the likelihood. Septal strain has only a minor reduction in uncertainty for the first three designs, yet using ***f***_4_ for parameter inference reduces septal strain uncertainty significantly. Moreover, using this final experimental design constrains S wall strain to have a similar shape to that of the LV and RV. Lastly, we quantify changes in mean pulmonary artery pressure, RV stroke volume, arterial and end-systolic ventricular elastance, and ventricular-vascular coupling for the different experimental designs. Histograms showing the frequency of these variables using the 600 forward samples are shown in Fig 11. Mean pulmonary artery pressure and arterial elastance have a comparable histogram width for all four experimental designs. In contrast, RV stroke volume, RV end systolic elastance, and ventricular vascular coupling have a larger variance in designs ***f***_1_ and ***f***_3_, which is reduced in designs ***f***_2_ and ***f***_4_.

**Fig 10.**
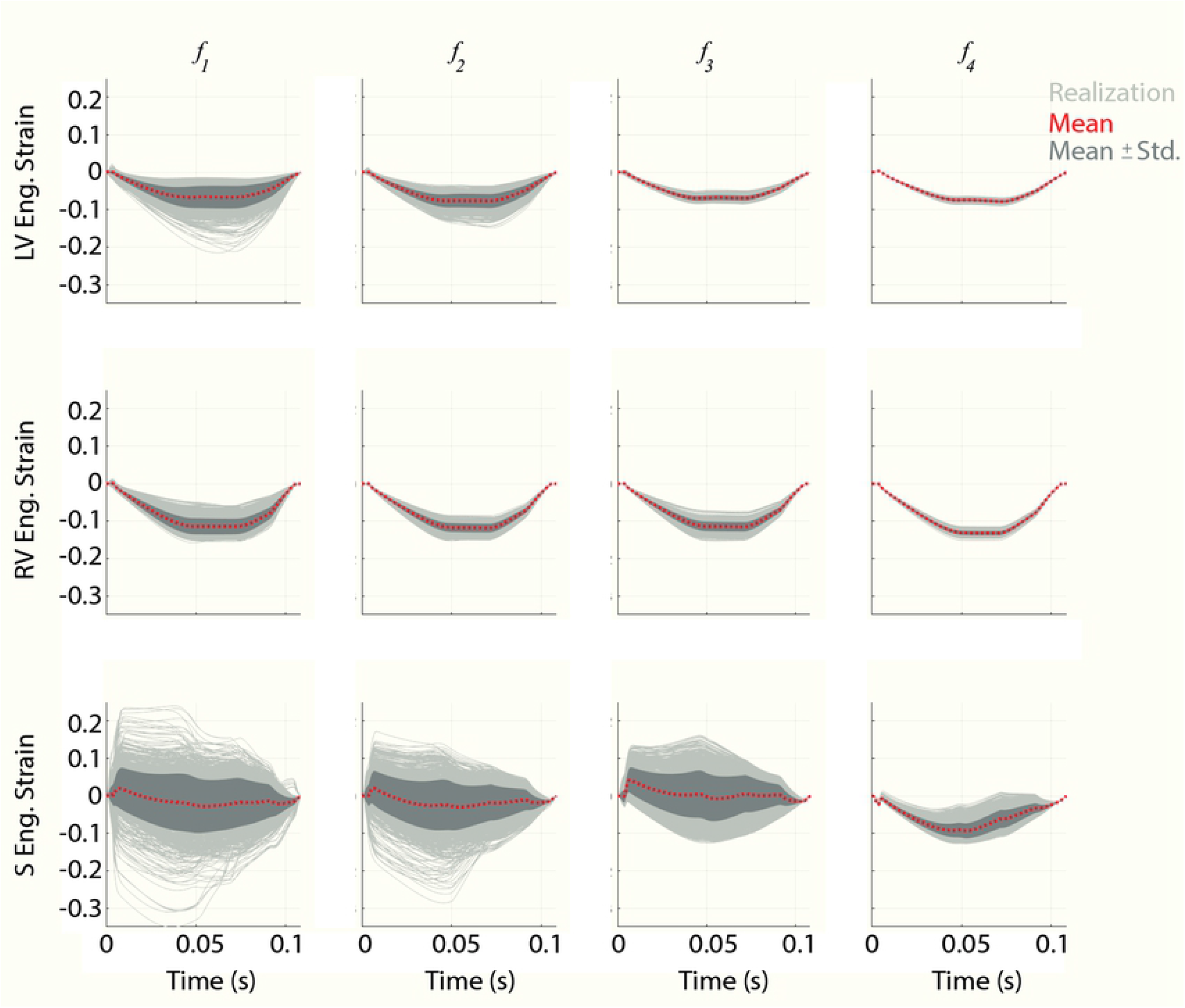
Forecast uncertainty in LV, RV, and S wall strain. Realizations in the LV, RV, and S engineering strain, along with the mean and ± standard derivation (Std), obtained from the posterior distributions. For designs only including RV dynamics (***f***_1_ and ***f***_2_), S engineering strain has a large uncertainty in the direction of strain (i.e., leftward or rightward). Designs including LV data (***f***_3_ and ***f***_4_) reduce the range of S strains, with the design ***f***_4_ ensuring that S strain is in the same direction as the LV. LV and RV strain have substantially less uncertainty than that of S, which shrinks with more informative designs.

**Fig 11.**
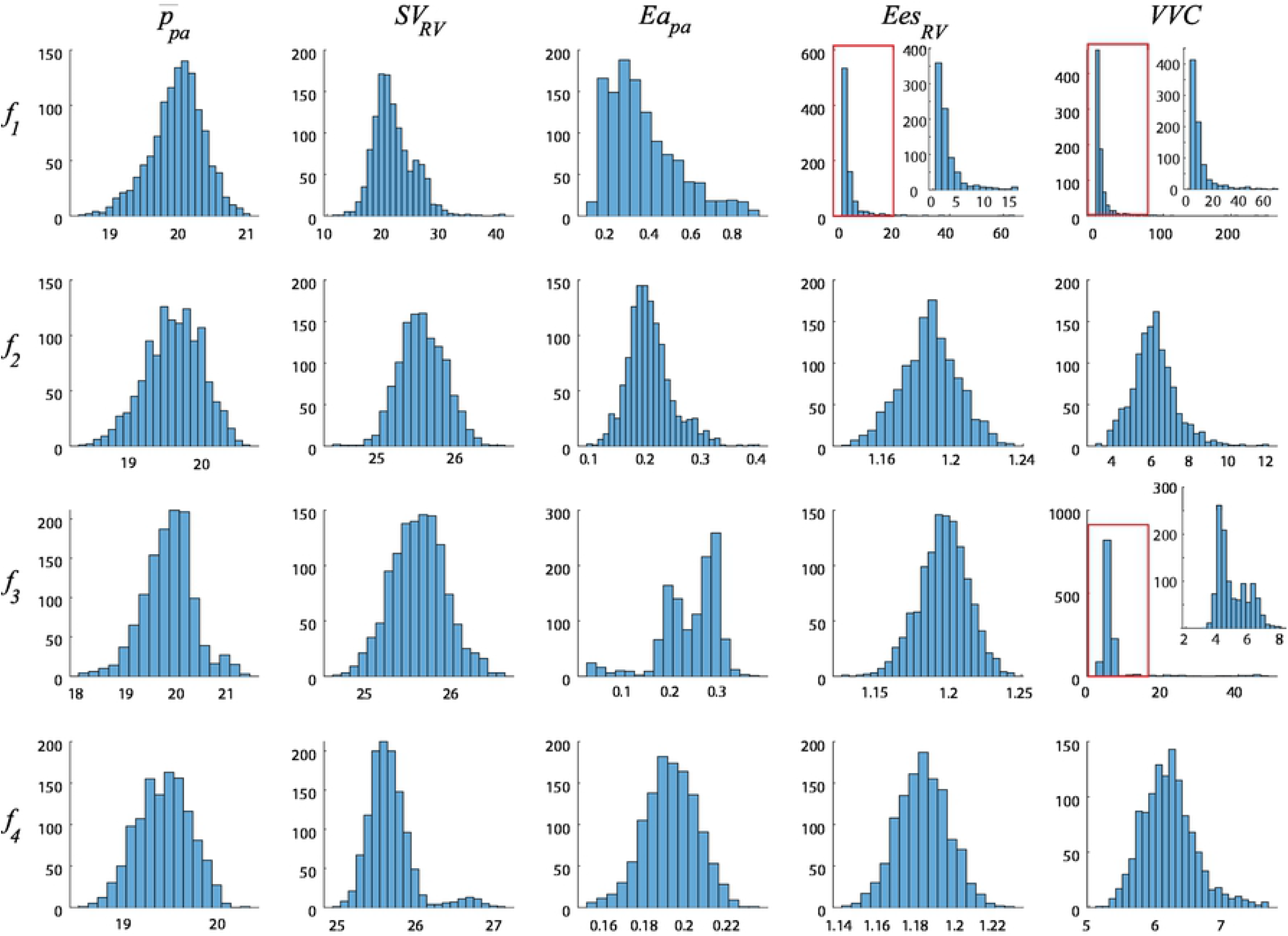
Simulated output quantities that are typically recorded when studying PH. Histogram plots of outputs typically recorded during *in-vivo* studies of PH progression are generated using the same 600 samples from the posterior that were used in Fig. 8, 9, and 10. These include mean pulmonary artery pressure 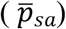, RV stroke volume (*SV_RV_,* defined as difference between maximum and minimum RV volumes), pulmonary arterial elastance (*Ea_pa_,* defined the difference between 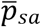 and mean LA pressure divided by *SV_RV_*), RV end systolic elastance (*Ees_RV_,* defined as the end systolic ratio of RV pressure and RV volume), and ventricular-vascular coupling (*VVC,* defined as the ratio *Ees_RV_/Ea_pa_*). Differences in the experimental design had little effect on 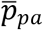. As expected, *SV_RV_* was more accurately captured with additional RV volume data. The wide variability in values of *Ea_pa_, Ees_RV_,* and *VVC* using the design ***f***_1_ is remedied once additional volume data is included in the design. Note that output values of *VVC* are made substantially more precise with additional LV data in ***f***_3_ and ***f***_4_.

## DISCUSSION

The present study investigates parameter identifiability for a multiscale model of cardiovascular dynamics. This work examines four different *in-vivo* experimental designs using *in-silico* modeling, and subsequently compares the reduction in parameter and output uncertainty under these different designs. *In-vivo* experimental designs are typically determined before using *in-silico* methods to analyze the data (9,30); however, some studies have considered using the latter to design optimal designs *a-priori* (44).

Sensitivity analyses are commonly used to reduce parameter sets to a smaller, more influential group (1,5,18). We use these techniques to reduce the original set of 49 parameters in the model to a set of 13 influential parameters. These 13 include those attributed to the TriSeg geometry, those describing the timing, duration, and active force of sarcomere shortening, and parameters describing the systemic and pulmonary vasculature, as shown in Fig 3. Similar to our analysis, the study by van Osta et al. (5) used Morris screening to calculate model sensitivity. They concluded that LV, RV, and S geometry parameters were most influential on simulations of chamber strain. While chamber strain was not considered as one of our output quantities of interest, our results suggest that these same parameters are influential on ventricular pressure and volume simulations. Their results also showed that, similar to our results, a single parameter from the left atrium was influential (5). We consider pressure and volume in the RV and LV as our outputs of interest, contributing to the addition of three influential circulatory parameters (*R_pul_, C_pa_*, and *R_sys_*). Both *R_pul_* and *C_pa_* describe the pulmonary vasculature, while only one systemic vasculature parameter, *R_sys_,* is influential and identifiable by our criteria. These three parameters were also influential in the analysis by Harrod et al. (1), who investigated PH due to LV diastolic dysfunction. The four experimental designs considered in this work focus on PH and RV function, hence more pulmonary parameters are influential than systemic. A majority of the influential parameters identified here are common in 0D models (1,18,30) and models incorporating the TriSeg framework (4,5,21), thus making the present analysis pertinent to future modeling studies utilizing either of these approaches.

Though local and global sensitivity analyses can identify influential parameters, they do not guarantee that parameters are identifiable (16). The sensitivity-based Fisher information matrix provides information about local parameter interdependence as well as quadratic approximations of parameter confidence intervals. In contrast, the profile likelihood analysis provides explicit confidence intervals for each parameter. The local sensitivity analysis performed here did not reveal identifiability issues, yet the profile likelihood based analysis illustrated limited identifiability across the less detailed experimental designs. This confounding result is documented in the review by Wieland et al. (16), suggesting again that profile likelihood analyses are superior in deducing parameter identifiability for nonlinear models. To the authors’ knowledge, the work by Pironet et al. (15) is the only other cardiovascular modeling study to consider this methodology. Their study integrated static pressure and volume data over multiple cycles, concluding that several parameters, including total stressed volume and vena cava compliance, were practically non-identifiable. The results in Fig 4 show that six of the parameters were identifiable across all four experimental designs. Of these, five describe the structure and function of the RV and pulmonary circuit, and the last describes systemic artery resistance. Interestingly, it appears that pulmonary vascular resistance, *R_pul_*, is practically non-identifiable using ***f***_1_, but should become identifiable for larger parameter bounds. This parameter is often used to describe PH and the state of the pulmonary vasculature, highlighting the need for additional data in the experimental design. Parameters describing the chamber wall volumes were consistently difficult to infer, especially in the LV and S when no LV data was available. This again suggests that a true understanding of heart function requires sufficient data on both ventricles.

This study is the first to compare multiple experimental designs using a multiscale model with biventricular interaction. As expected, increasing the amount of data available in the likelihood function reduced the confidence interval width, i.e., more data decreases the uncertainty in the estimates. In contrast to prior studies utilizing the profile likelihood (15), our results show spikes in the likelihood values. We accredit this non-smoothness to possible incompatibilities in the DAE system, which can frequently occur with this model (5,19), returning large values in the residual sum of squares. However, we are primarily interested in using the profile likelihood method to test for whether the confidence intervals have finite bounds; hence, smooth profile likelihoods are not necessary to determine if the parameter subsets are identifiable. Overall, the most complete experimental design, *f*_4_, led to the tightest confidence intervals and reduced numerical instabilities, though perturbations in *V_wall,LV_, V_wall,RV_*, and *V_wall,S_* still cause some sharp jumps in the likelihood.

MCMC is a common tool for Bayesian inference, and can also assess parameter uncertainty and identifiability (1,18,19,39). The posterior densities in Fig 5, Fig 6, and Fig 7 suggest that most of the parameters are practically identifiable in the presence of measurement noise. The wall volumes, *V_wall_*, have wider posteriors across the first two experimental designs, but tend to shrink with additional LV data, consistent with the profile likelihood results. Specifically, *V_wall,LV_* has a nearly uniform posterior when only using RV pressure data (***f***_1_), suggesting practical identifiability issues. This is expected, as this parameter has its largest effects on LV dynamics, which are only present in designs ***f***_3_, and ***f***_4_. The active stress parameter, *σ_act,v_,* is not identifiable using ***f***_1_, and has a long tail towards larger values. This parameter shows noticeable changes in mixing properties when using the systolic and diastolic LV outputs in ***f***_3_, and may be due to sampling in higher rejection regions to obtain appropriate LV outputs. All twelve pairwise plots in S2 text using the design ***f***_4_ show somewhat strong correlations between *V_wall,LV_* and *σ_act,v_,* as well *A_m,ref,LA_* and *C_pa_.* Contrasted with the profile likelihood results, this may suggest that these parameters are not practically identifiable given measurement noise. The posteriors using ***f***_3_ and ***f***_4_ in Fig 5, Fig 6, and Fig 7 are nearly all unimodal, with the true data generating parameters located near the modes. As noted by Paun et al. (39), flat, uniform posteriors suggest that parameters are not practically identifiable, supporting our claim of improved identifiability with more detailed experimental designs. The study by Harrod et al. (1) also used MCMC to test for structural identifiability; however, their results show a deviation between the true value of *R_sys_* and the posterior distribution, whereas our results (for ***f***_2_, ***f***_3_, and ***f***_4_) show an overlap in the true and estimated values. Discrepancies between Harrod et al. and our results are likely attributed to differences in model complexity and data availability for parameter inference. van Osta et al. (19) constructed parameter posteriors for *A_m,ref_* and the equivalent of our *σ_act,v_* and *τ_sys,v_* using MCMC. Their study also showed that repeated construction of the posteriors from different initial guesses had reasonable overlap, suggesting all parameters were identifiability. Colunga et al. (18) contrasted two parameter subsets using MCMC and heart-transplant data. The non-identifiable set had posteriors with long, unbounded tails, whereas, like the results here, the identifiable set has tighter posterior distributions with finite tails. The posterior variance (the width of the posteriors) shown in Fig 5, Fig 6, and Fig 7 tends to decrease with increased data availability, consistent with the profile likelihood confidence intervals. A comparison of the hemodynamic posteriors in Fig 7 reveals that both *R_pul_* and *R_sys_* have larger uncertainty when only using RV pressure in the design ***f***_1_. As noted previously in the text, pulmonary vascular resistance is a pertinent biomarker of PH progression and severity (23,45). Our results suggest that, at a minimum, RV volumes are included in the experimental design to obtain reasonable estimates of hemodynamic parameters and better constrain posterior widths for *V_wall_* parameters. Interestingly, both the profile likelihood analysis and the MCMC results suggest that *C_pa_* is identifiable but without an improvement with more complex designs. This may be attributed to the simplicity of the pulmonary artery compartment, and may vary more if using a more complex model of the proximal pulmonary arteries (2).

Whereas the parameter posteriors provide insight into their uncertainty, sampling from these posteriors describes uncertainty in the model output. The first design, ***f***_1_, provides information about RV pressure, and corresponding model simulations shown in Fig 8 have little uncertainty. In contrast, *V_RV_*(*t*),*p_LV_*(*t*), and *V_LV_*(*t*) exhibit larger uncertainty, as shown in Fig 8 and Fig 9, with the mean response often deviating from the true signal. The more data-rich experimental designs lead to a better agreement between the model and the simulated data as well as a reduction in uncertainty. Interestingly, differences in uncertainty bounds between ***f***_3_ and ***f***_4_ are not evident in the isolated pressure and volume signals in Fig 8, yet PV loop uncertainty in the LV is reduced substantially in Fig 9. The difference in these two plots is linked to the timing of ventricular PV dynamics, which become more apparent when plotting pressure versus volume. The reduction in uncertainty when the design ***f***_3_ is used suggests that including static systolic and diastolic measures of LV function are sufficient for model calibration and are necessary to reduce output uncertainty. This experimental design was utilized by Philip et al. (23) in a mouse model of PH due to left heart failure. Their results highlighted that impaired LV function can ultimately raise pulmonary vascular resistance and contribute to RV dysfunction. Assessing the LV via echocardiography is easier than the RV due to anatomic shape and location (46), hence adding this assessment to dynamic RV PV loop protocols is reasonable and provides insight into LV impairment during PH (11). Recent studies have also found significant changes in both left and right atrial function in heart failure and PH (23,47,48). We found only one atrial parameter, *A_m,ref,LA_,* was influential and identifiable on RV and LV outputs. Allowing this parameter to vary explains the greater variability in left atrial PV loops than the corresponding right atrial simulations in the first three designs. However, it seems that dynamic data in the LV reduces the variability in left atrial forecasts, suggesting that ***f***_4_ is the most optimal design for studying left atrial function in the absence of left atrial data. We did not consider atrial data in our possible designs, yet future work may reveal its significance in understanding disease progression, especially PH due to left heart failure (1,23).

The TriSeg model is an efficient simulator of biventricular interaction. Prior work has used this model to quantify changes in biventricular interaction under diseases such as PH (21,49), arrhythmogenic cardiomyopathy (19), and mechanical desynchrony (50). Our results in Fig 10 show that the uncertainty in LV, RV, and S wall strain tend to decrease with more informed experimental designs. Though the model employed van Osta et al. (19) has fundamental differences from our model, both have comparable uncertainty in wall strains. Their study calibrated model predictions to measurements of wall strain by echocardiography, yet our work shows that calibration to pressure and volume data is sufficient in reducing simulated wall strain uncertainty. Strain forecasts also elucidate the state of LV-RV interaction, which is compromised in the presence of PH (11).

We use the model to simulate other hemodynamic quantities typically recorded in PH studies (23). The distribution of simulated mean pulmonary arterial pressure in Fig 11 are similar in width across the experimental designs. Both *R_pul_* and *C_pa_* play a role in this output, yet Fig 7 shows that *R_pul_* has a noticeably smaller posterior when informed by ***f***_4_. Though *R_pul_* will ultimately dictate the pressure magnitude, the unchanged posterior in *C_pa_* suggests that this parameter is largely attributed to mean pulmonary artery pressure. The study by Colunga et al. (18) found that including *R_pul_* and *C_pa_* in parameter inference led to close agreement between model predictions of pulmonary artery pressure and measured data. The uncertainty in mean pulmonary artery pressure described by Harrod et al. (1) are similar to our results as well. As expected, forecasts of RV stroke volume and pulmonary artery elastance (defined as the difference between mean pulmonary artery pressure and mean left atrial pressure divided by the RV stroke volume) have small variability with any designs including RV volume, i.e., ***f***_2_, ***f***_3_, and ***f***_4_. Hence, the relatively wide probability densities for RV end-systolic elastance and RV ventricular-vascular coupling are directly tied to uncertain model predictions of RV volume. A zoom of the model forecasts shown in Fig 11 shows that additional volume constraints narrow the output uncertainty in these indices. All five indices examined here can be indicative of PH progression and RV function and suggest that RV pressure alone is not informative enough to constrain the model forecasts. Therefore, future experiments into PH and RV function should strive to have both RV pressure and volume data collected, along with static or dynamic measures of LV function.

There are several limitations in this study. We generate synthetic data from our mathematical model to test for identifiability, hence our noise model correctly matches the true added noise. When using physiological data, this may not hold true, and may require additional components to the statistical model (e.g., model discrepancy (39)). The profile likelihood results presented here exhibit non-smoothness, whereas prior studies (17,51) typically show smooth profiles. This could be obtained by considering more sophisticated parameter mesh refinement, as the system of DAEs are stiff and can lead to model failure if parameters are not compatible in the model. We consider four experimental designs that are typically used to assess RV function, yet numerous other designs including additional data (e.g., systemic artery pressure, pulmonary artery wedge pressure) could provide unique insight into RV function. Our analysis is applied to data simulated for a normotensive mouse as opposed to simulating PH data. However, we believe the present analysis will be consistent even when parameters are adjusted to the PH range.

## CONCLUSION

The present study investigates parameter identifiability of a cardiovascular model with biventricular interaction, specifically calibrated for mouse hemodynamics. Using a combination of sensitivity analyses, profile likelihood confidence intervals, and MCMC, we construct a subset of 13 influential parameters. The present analyses are conducted on model outputs corresponding to four experimental designs used to study PH and RV failure *in-vivo*. Profile likelihood analysis shows that model parameters are not uniquely identifiable when only RV pressure data is available, and that more informed designs are necessary to recapture the true parameter values. Using noise corrupted data, we show that parameter posteriors are modulated with more informative designs, and that model parameters are more precisely identified with the addition of LV data. Model forecasts of LV dynamics are more uncertain when designs do not account for LV pressure and volume, whereas more informed experimental designs reduce forecast uncertainty in both PV loops and wall strain. We conclude that future, synergistic studies using both *in-vivo* and *in-silico* methods should incorporate functional LV data to improve model forecasts of cardiac function and biventricular dynamics. We hypothesize that this will be especially important when studying the progression of RV failure due to PH.

## CITATION DIVERSITY STATEMENT

In agreement with the editorial from the Biomedical Engineering Society (BMES) (52) on biases in citation practices, we have performed an analysis of the gender and race of our bibliography. This was done manually, though automatic probabilistic tools exist (53). We recognize existing race and gender biases in citation practices and promote the use of diversity statements like this for encouraging fair gender and racial author inclusion.

Our references contain 9.25% woman(first)/woman(last), 14.8% man/woman, 16.7% woman/man, and 59.3% man/man. This binary gender categorization is limited in that it cannot account for intersex, non-binary, or transgender people. In addition, our references contain 3.70% author of color (first)/author of color(last), 5.55% white author/author of color, 25.9% author of color/white author, and 64.8% white author/white author. Or approach to gender and race categorization is limited in that gender and race are assigned by us based on publicly available information and online media. We look forward to future databases that would allow all authors to self-identify race and gender in appropriately anonymized and searchable fashion and new research that enables and supports equitable practices in science.

## APPENDIX

### Sarcomere model

We consider the sarcomere model based on prior work (8,27). The sarcomere is modeled as two series passive elements (describing extracellular matrix (ECM) and Titin contributions) in parallel with a series combination of an active, contractile element and a series, elastic element. Both the ventricles and atria are modeled using identical formulations and deviate only in their parameter values.

The sarcomere length is calculated as a function of the myocardial strain

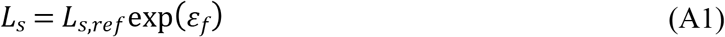

where *L_s,ref_* = 2.0*μ*m is the reference sarcomere length at zero strain (i.e., *ε_f_* = 0). The exponential function mimics the nonlinear behavior of sarcomere length under load. Assuming that the change in contractile element length, *L_sc_,* depends linearly on the change in sarcomere length, the dynamics of *L_sc_* (μm) are

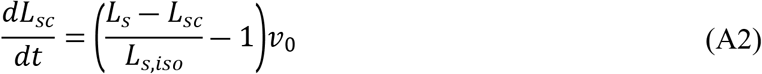

where *L_s,iso_* (*μ*m) is the length of the elastic series element in an isometrically stressed state, and *v*_0_ (*μ*m/s) is the velocity of sarcomere shortening with zero load (8).

The mechanical activation of the sarcomere is heuristically modeled as separate “rise” and “decay” functions. The former is given by

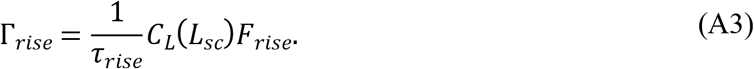

*C_L_* and *F_rise_* represent the increase in contractility with sarcomere length and with time, respectively. They are given by

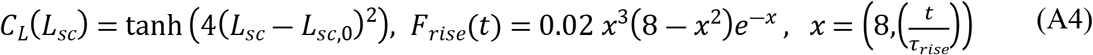

where *τ_rise_* (s) scales the rise in contractility and *L*_*sc*,0_ (*μ*m) represents the contractile element length with zero active stress. The decay function is

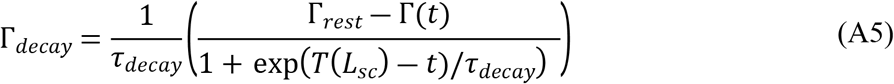

where Γ_*rest*_ is the diastolic resting level of activation, *τ_decay_* (s) is the decay time, and

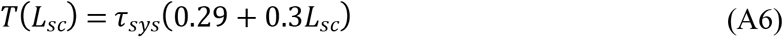

represents the decay in activation with decreasing sarcomere length, which depends on the duration of systole, *τ_sys_* (s). Both rise and decay functions are combined to represent the change in contractility

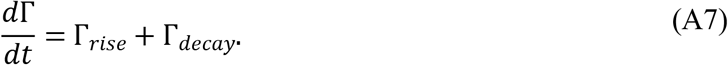

The active stress, *G_act_* is finally calculated as the product of contractility, contractile element displacement, and the strain of the series elastic element

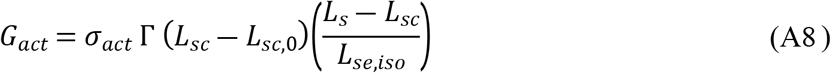

where *σ_act_* (KPa) is a parameter that scales the active force contribution.

For the passive stress, we consider a new, *passive* reference length *L_s,pas,ref_* (*μ*m), giving the passive strain

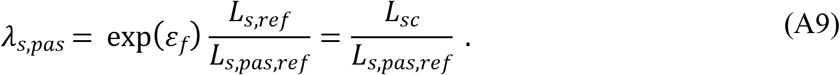

The two passive stresses are as defined in Walmsley et al. (54)

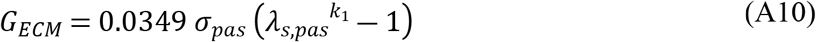

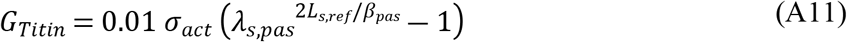

where *σ_pas_* (KPa) is the factor scaling the passive force of the ECM, *k*_1_ (dimensionless) is an exponential scaling parameter, and *β_pas_* (*μ*m^-1^) is a measure of Titin stiffness (54).

### Cardiac chamber equations

For the atria, the mid-wall volume, *V_m_*, mid-wall curvature, *C_m_*, and mid-wall cross-sectional area, *A_m_,* are determined by

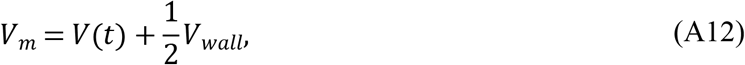

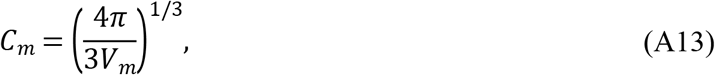

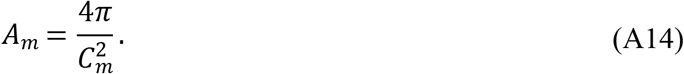

Since the LV, RV, and S are mechanically coupled, a separate formulation for *V_m_, C_m_,* and *A_m_* is required. Utilizing the common radius of mid-wall junction point *y_m_* and denoting the maximal axial distance from each chamber wall surface to the origin as *x_m_* (8,29), we get

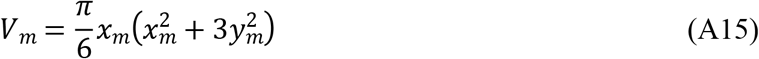

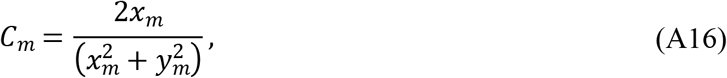

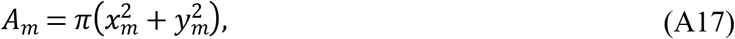

for the LV, RV, and S. As in the original work by Lumens (8), we can also related *V_m_* to the blood volume V(t) in the chamber. Specifically, *V_m,LV_* and *V_m,RV_* are

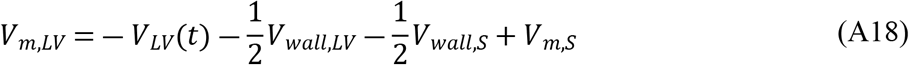

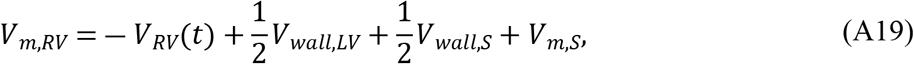

which is updated at each time point. Hence, we solve for *x_m_* using in equation (5) to deduce *A_m_* and *C_m_* for each TriSeg component. Finally, the atrial transmural pressure is determined from the wall tension and mid-wall curvature

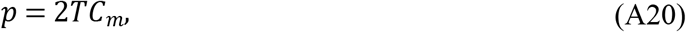

while the ventricular pressure in the TriSeg model is calculated by (8)

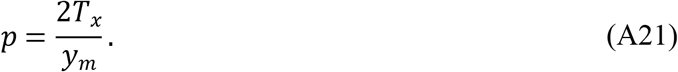

